# Microglial SIRT2 deficiency aggravates cognitive decline and amyloid pathology in Alzheimer’s disease

**DOI:** 10.1101/2025.01.17.633523

**Authors:** Noemi Sola-Sevilla, Maider Garmendia-Berges, Mikel Aleixo, MCarmen Mera-Delgado, Maite Solas, Rosa M Tordera, Lucía García-Carracedo, Sara Expósito, Elena Anaya-Cubero, Joaquín Fernández-Irigoyen, Elisabeth Guruceaga, Enrique Santamaria, Eduardo D. Martin, Elena Puerta

## Abstract

Sirtuin 2 (SIRT2), a NAD+-dependent deacetylase, has been implicated in aging and neurodegenerative diseases such as Alzheimer’s disease (AD). While global SIRT2 inhibition has shown promise in reducing amyloid-beta pathology and cognitive deficits in different mouse models of AD, peripheral SIRT2 inhibition has been associated with adverse effects, such as increased inflammation. This suggests that targeted inhibition of specific cellular populations within the brain may represent a more precise and effective approach for the treatment of AD. To explore this hypothesis, we generated a conditional microglial SIRT2 knockout mouse model in the context of AD. Our results reveal that microglial SIRT2 reduction does not confer protective effects in the APP/PS1 model; rather, it aggravates cognitive decline, accelerates amyloid plaque deposition, and increases levels of pro-inflammatory cytokines at early stages of AD pathology. Transcriptomic analysis further indicates that SIRT2-deficient microglia exhibit altered expression of genes associated with aging and synaptic dysfunction. This phenotype was accompanied by increased phagocytosis of synaptic elements and impaired long-term potentiation. These findings suggest that while SIRT2 inhibition in some contexts may be beneficial, targeted inhibition within microglia could accelerate AD progression, underscoring the need for cell-specific approaches when considering SIRT2 as a therapeutic target.

## 1. Introduction

Sirtuins, a family of highly conserved NAD+-dependent histone deacetylases, exert significant influence over numerous biological and pathological processes. Specifically, the intricate interplay between sirtuins and neurodegenerative diseases has gained significant attention in recent years [1]. Among these sirtuins, sirtuin 2 (SIRT2) stands out as a pivotal player, particularly in the central nervous system (CNS), where its abundance [2] underscores its potential significance in aging and age-related pathologies.

However, the dichotomy between its roles in the CNS and the periphery presents a complex landscape that demands meticulous investigation (for a review see [3]). On the one hand, aging correlates with heightened SIRT2 expression in the brain [4–7], suggesting the potential interest of its pharmacological inhibition as a therapeutic strategy for neurodegenerative diseases. Accordingly, SIRT2 inhibition or its genetic deletion has provided beneficial effects in different mouse models of Huntington’s disease [8,9], Parkinson’s disease [10–12], and Alzheimer’s disease (AD) [13–16]. Additionally, it has been also proposed as an effective pharmacological option for the treatment of neuropsychiatric disorders such as depression [17–19]. On the other hand, the analysis of the peripheral consequences of SIRT2 inhibition or deletion presents a contrasting storyline. Here, decreased SIRT2 levels with age have been associated with age-related inflammation, insulin resistance, and cardiovascular diseases [20–23]. Thus, systemic administration of modulators of this enzyme to treat neurodegenerative diseases would have detrimental outcomes that should not be underestimated. In fact, a recent study published by our group has demonstrated that systemic SIRT2 pharmacological inhibition improves cognitive dysfunction and reduces amyloid pathology and neuroinflammation in a transgenic mouse model of AD [14]. However, the same treatment increases peripheral levels of pro-inflammatory cytokines (IL-1β, TNF, IL-6 and MCP-1) recapitulating the pathologic phenotype observed in old SIRT2 knockout mice [20] and supporting the notion that peripheral SIRT2 is essential to prevent aging related inflammation. These disparities emphasize the need for a deeper comprehension of the distinct roles played by SIRT2 in various cell types, which is crucial for developing novel approaches to target SIRT2 in therapeutic interventions.

AD stands as the most prevalent form of dementia, with a staggering projection of nearly 131 million cases by 2050. The complex etiopathology of AD encompasses a multitude of pathological factors, including neuroinflammation and amyloidogenic proteins such as amyloid-β (Aβ). Notably, genome-wide association studies in AD have unveiled that a significant proportion of genes identified at risk loci are intricately linked to neuroinflammation and predominantly expressed in microglia, the main resident immune cells of the CNS (for a comprehensive review, refer to [24]). These findings support the notion that neuroinflammation plays a key role on both pathogenesis and progression of AD [25–28] challenging previous assumptions that it solely manifests as downstream effect or consequence of primary pathological lesions. In fact, in recent years, microglia have been increasingly recognized as key modulators of AD pathogenesis [29–31]. These cells are actively involved in synaptic plasticity, where they influence synapse formation, function, and elimination during both development and adulthood [32]. Additionally, microglia are specialized in sensing and responding to pathogens, misfolded proteins, or molecules released by damaged cells. The balance between the release of cytotoxic and neurotrophic molecules, along with the timing and the intensity of these responses is crucial in determining whether microglial activity will be beneficial or detrimental to the CNS [33]. Therefore, deciphering the mechanisms that regulate microglial functions is essential for developing therapeutic strategies for neurological diseases.

Taking into account the contrasting conclusions obtained when SIRT2 is inhibited in the CNS or at the peripheral level, in the present study we hypothesize that targeting SIRT2 inhibition specifically in CNS microglia is the best approach. Thus, by elucidating the impact of microglial SIRT2 deficiency on AD pathology, we aim to delineate the potential therapeutic benefits while circumventing the adverse systemic effects associated with global SIRT2 inhibition.

## 2. Material and Methods

### 2.1. Animals

To generate the microglial conditional SIRT2 knockout mouse model, C57BL/6-Tmem119^em1(cre/ERT2)Gfng/J^ (Strain number 031820) and B6.129(Cg)-Sirt2^tm1.1Auw/J^ (Strain number: 030835) mice were purchased from Jackson Laboratory. These two strains were bred for several generations to obtain the Sirt2^flx/flx^-Tmem119^cre/ERT2^ mouse model. To evaluate the behavioral and molecular consequences of microglial SIRT2 deletion on the etiology and the progression of AD, APPswe-PSEN1dE9 (named APP/PS1) transgenic mice on a C57BL/6;C3H genetic background (MMRRC Strain number: 034829-JAX; Jackson Laboratory, Bar Harbor, ME, USA) were bred with Sirt2^flx/flx^- Tmem119^cre/ERT2^ mice to obtain the APPswe-PSEN1dE9: Sirt2^flx/flx^-Tmem119^cre/ERT2^ mouse model (named APP/PS1-SIRT2^ΔTmem119^). The APPswe-PSEN1dE9 and Tmem119-CreERT2 alleles were maintained in heterozygous conditions, whereas the floxed SIRT2 allele was maintained in homozygous conditions. Since microglia have been shown to regulate neurogenesis and synaptic maturation at perinatal and postnatal stages [34], to achieve the cre-dependent deletion of *Sirt2* gene, tamoxifen (100 mg/kg i.p. every 24 h for 5 days) was administered when mice were 6 and 10 weeks old. The significant reduction of *Sirt2* expression was confirmed in isolated microglia from the hippocampus of 5 month-old animals (Figure S1). 5 and 8-month-old male and female control, SIRT2^ΔTmem119^, APP/PS1 and APP/PS1-SIRT2^ΔTmem119^ littermates were used for the experiments.

Animals were housed in groups in standard breeding cages (5 mice per cage) and had access to food and water *ad libitum*. Temperature and humidity were constant (23 ± 1 °C and 55 ± 10%, respectively), and lights were maintained on a 12 h light/dark cycle (light-dark: 8:00AM–8:00PM). All efforts were made to minimize animal suffering and reduce the number of animals used in the experiments. The procedures were conducted in accordance with the principles of laboratory animal care outlined in the European Communities Council Directive 2013/53/EC and reported in compliance with the ARRIVE guidelines. Ethical approval was obtained from the University of Navarra’s ethics committee.

### 2.2. Behavioral tests

All behavioral tests were monitored by a video camera connected to the *Ethovision XT 11.5* tracking system (Noldus, Wageningen, The Netherlands).

#### Spontaneous motor activity

Motor activity was measured for 15 min in an open field (35 x 35 cm^2^). The experimental room was softly illuminated, and each mouse was placed in one cage. Distance moved and speed were recorded for each animal.

#### Elevated plus maze

Elevated plus maze (EPM) is an elevated platform (38.5 cm above the ground) with two close and two open arms (30 x 5 cm^2^ each), crossed in the center oppositely one to another with a middle region of 5 x 5 cm^2^. Mice were placed in the center of the maze and were permitted to move freely between them for 5 min. The frequency to enter in the open arms and the total time spent in them were used as a measure of an anxiety-like behavior.

#### Marble burying test

Fifteen glass marbles were evenly distributed in a cage (3 marbles per row, 5 rows), and a mouse was placed in the center of the cage. After 30 minutes, the number of buried marbles was recorded as a measure of the mouse’s anxious behavior.

#### Forced swim test

For forced swim test (FST), glass cylinders (21 cm height, 13 cm diameter) filled with 15 cm of water maintained at 23 ± 1 °C were used. Mice were placed individually into each glass for 6 min and the duration of immobility was recorded during the last 4 min of the testing period. Immobility was calculated as the time when the mouse made only small movements to keep its head above the water.

#### Morris water maze

The evaluation of spatial memory and working and reference memory was performed by using the Morris water maze (MWM). The water maze consisted of a circular pool with a diameter of 120 cm filled with water (21-22 °C) and virtually divided into four equal quadrants (northeast, northwest, southeast, and southwest). To guide the mice, visual cues were strategically positioned both in the room and inside the walls of the pool.

During the visible-platform training phase (2 days, 6 trials per day), a platform was situated in the southwest quadrant raised above the water with an object placed on top to facilitate its location (Habituation phase). To evaluate learning capacity (Acquisition phase), a hidden platform (1 cm below the water surface) was placed in the northeast quadrant of the pool. The trial concluded when the animal reached the platform (escape latency) or after 60 s in the pool. If the mouse did not find the platform within 60 s, it was guided to the platform and allowed to stay there for 15 s. The test was conducted over 7 consecutive days (4 trials per day). For memory retention assessment, the platform was removed from the pool, and animals were allowed to swim for 60 s (Retention phase). This trial took place on the 5^th^ and 8^th^ days (last day) and the percentage of time spent in the northeast quadrant was recorded. All trials were monitored by a video camera set above the center of the pool.

### 2.3. Electrophysiology

Synaptic transmission in hippocampal slices was analyzed as previously described [35]. Briefly, transverse brain slices of 350 μm thick were cut with a vibratome (VT1200S, Leica Microsystems, Wetzlar, Germany). Then, they were incubated for at least 1 h at RT in artificial cerebrospinal fluid (aCSF) that contained 124 mM NaCl, 2.69 mM KCl, 1.25 mM KH_2_PO_4_, 2 mM MgSO_4_, 26 mM NaHCO_3_, 2 mM CaCl_2_ and 10 mM glucose, gassed with a 95% O_2_/5% CO_2_ mixture at pH 7.3–7.4. Individual slices were then transferred to an immersion recording chamber and perfused with oxygenated warmed aCSF (32 ± 2 °C). Field excitatory postsynaptic potentials (fEPSPs) were recorded in the *stratum radiatum* of CA1 pyramidal layer by a carbon fiber microelectrode (Carbostar-1, Kation Scientific, Minneapolis, MN, USA). Evoked fEPSPs were elicited by stimulation of the Schaffer collateral fibers with an extracellular bipolar tungsten electrode placed in the *stratum radiatum* via a 2100 isolated pulse stimulator (A-M Systems, Inc., Carlsborg, WA, USA). At the beginning of each experiment, basal synaptic transmission was analyzed by applying isolating stimuli of increasing intensity to reach a maximal fEPSP response. For long-term potentiation (LTP) experiments, the stimulus intensity was adjusted to elicit 50% of the maximum response signal and kept constant throughout the experiment. After recording stable baseline responses for 30 min, LTP was induced by a single train of theta burst stimulation (5 bursts of 5 pulses at 100 Hz, with an interval of 200 ms between bursts). Potentiation was measured for 1 h after LTP induction at 0.2 Hz.

### 2.4. Bulk-RNA sequencing

Mice were transcardially perfused with ice-cold PBS for 6 minutes under anesthesia with xylazine (10 mg/kg) and ketamine (80 mg/kg). Following perfusion, the brains were removed, and the hippocampi were dissected on ice. The samples were digested at 37 °C with rotation for 30 min with papain (2 mg/mL, Worthington Biochemical Corporation, Lakewood, NJ, USA) in Dulbecco’s PBS (Lonza, Basel, Switzerland), containing 50 μg/mL of DNase I (cat# 11284932001; Hoffmann-La Roche, Basel, Switzerland), 5 µg/mL of actinomycin D (cat# A1410), 10 µM of triptolide (cat# T3652), and 27.1 µg/mL of anisomycin (cat# A9789) (all from Sigma-Aldrich, Saint Louis, MO, USA). Then, the tissue was mechanically dissociated and filtered through a 70 μm cup Filcon cell suspension filter (BD, Franklin Lakes, NJ, USA), followed by a centrifugation at 300 rcf for 15 min at 4 °C. For myelin removal, the cellular pellet was resuspended in 1 mL of 25% Percoll solution (GE Healthcare Bio-Sciences AB, Chicago, IL, USA) and centrifuged at 1000 rcf for 10 min at RT. Cell suspensions were incubated with CD11b MicroBeads at 1:10 dilution (Miltenyi Biotec, Bergisch Gladbach, Germany) for 15 min at 4 °C. After washing the cells with autoMACS buffer (0.5% FBS, 1% EDTA and 1% penicillin/streptomycin in PBS 1x), Microglial CD11b^+^ cells were separated on an *autoMACS Pro Separator* (programs: *Posseld* and *Rinse*; Miltenyi Biotec). RNA sequencing was performed by adapting the technology of SCRB-Seq [36] to allow for the high cost-efficient multiplexed transcriptome characterization. Briefly, cells were collected in cell lysis buffer and poly-(A)+ RNA were purified using the *Dynabeads mRNA DIRECT Purification Kit* (ThermoFisher Scientific) from approx. 10000 cells. poly-(A)+ RNA were annealed to a custom primer containing a poly-(T) tract, a Unique Molecule Identifier (UMI), and a sample barcode. Retrotranscription using Template-switching oligonucleotides (TSO) was then used to synthetize and amplify 3’UTR enriched cDNA, resulting in barcoded cDNA fragments. Library preparation was then completed using the *Nextera XT* (Illumina) protocol which introduces i5-P5 and i7-P7 structure for massive parallel sequencing. Quality control was performed following pre-amplification RT and library preparation to ensure quality and length accuracy, as well as to equilibrate sample pooling. Libraries were then sequenced using a NextSeq2000 sequencer (Illumina). 20 million pair-end reads (Rd1:26; Rd2:80) were sequenced for each sample and demultiplexed using bcl2fastq.

RNA sequencing data analysis was performed using the following workflow: (1) the quality of the samples was verified using *FastQC* software (https://www.bioinformatics.babraham.ac.uk/projects/fastqc/) and the trimming of the reads with trimmomatic [37]; (2) alignment against the mouse reference genome (GRCm39) was performed using STAR [38]; (3) gene expression quantification using read counts of exonic gene regions was carried out with *featureCounts* [39]; (4) the gene annotation reference was Gencode vM33 [40]; and (5) differential expression statistical analysis was performed using R/Bioconductor [41]. Data are publicly available in GEO database with the accession number GSE280235.

First, gene expression data was normalized with *edgeR* [42] and *voom* [43]. After quality assessment and outlier detection using R/Bioconductor [41], a filtering process was performed. Genes with read counts lower than 6 in more than 50% of the samples of all the studied conditions were considered as not expressed in the experiment under study. *LIMMA* [43] was used to identify the genes with significant differential expression between experimental conditions. Further functional and clustering analyses and graphical representations were performed using R/Bioconductor [41] and clusterProfiler [44]. Metascape was also used to extract biological information associated with transcriptomic functionality using default settings (minimum overlap: 3; minimum enrichment: 1.5; P < 0.01) [45].

### 2.5. Quantitative real-time PCR

Total RNA was isolated from cortex samples using TRI Reagent^®^ (Sigma-Aldrich). RNA was retro-transcribed into cDNA using the *High-Capacity cDNA Reverse Transcription Kit* (Applied Biosystems, Foster City, CA, USA). Quantitative real-time PCR (qPCR) was carried out using Taqman^®^ Universal PCR Master Mix (Applied Biosystems) on *ViiA™ 7 Real-Time PCR System* (Applied Biosystems), and *Gapdh* gene was used as internal control. The primers were purchased from Applied Biosystems (*Il-1β* cat# Mm00434228_m1; *Il-6* cat# Mm00446190_m1; *Tnf-α* cat# Mm00443258_m1; *Gapdh* cat# Mm99999915_g1).

For gene expression quantification, the double delta CT (ΔΔCT) method was used where delta CT (ΔCT) values represent normalized target genes levels with respect the internal control. The relative quantification of all targets was carried out using the comparative cycle threshold method, 2^−ΔΔCt^, where ΔΔCt = (Ct target gene − Ct endogenous control) treated / (Ct target gene − Ct endogenous control) untreated.

### 2.6. Dendritic spines morphology

*FD Rapid GolgiStain™ Kit* (FD NeuroTechnologies, Columbia, MD, USA) was used for the study of the morphology of dendritic spines. Briefly, after dissection, one brain hemisphere was rinsed in Milli-Q water and immersed in solution A/B previously prepared (see manufacturer’s protocol). The brains were stored in this solution at RT for 2 weeks protected from light. Then, brains were transferred into solution C and stored at RT protected from light. After 72 h, tissues were frozen with dry ice and stored at-80 °C.

Serial coronal brain slices (thickness: 90 μm) were cut with a *cryostat HM 500 OM* (MICROM International GmbH, Walldorf, Germany) at-20 °C and sections were mounted on gelatin-coated microscope slides (cat# PO101; FD NeuroTechnologies). Manufacturer’s instructions were followed for the staining procedure with solution D/E and the slices were mounted with *Eukitt^®^ Quick-hardening Mounting Medium* (Sigma-Aldrich). Images were acquired with the *Zeiss Axio Imager M1Esc* microscope and the program *ZEN 2 blue edition* (Zeiss Microscopy, Jena, Germany). The analysis was performed on apical collateral dendrites (stratum radiatum) from CA1 pyramidal neurons. A stack of 60-80 images (with 1 μm thick Z stacks) per dendrite were captured randomly based on the following criteria: (i) neurons relatively complete (at least two orders of dendrites entirely visible) and (ii) no overlap with other labeled neurons. This was performed on a total of 25 dendrites per animal (n = 4 animals per group) utilizing a 100x objective-equipped automated microscope (Axio Imager M1; Carl Zeiss Microscopy). For spine analysis and classification, *Reconstruct* software (version 1.1.0.0, 2007; http://synapses.clm.utexas.edu) was used following the method developed by [46] which facilitates impartial classification of spine types. All analyses were conducted blinded.

### 2.7. Immunofluorescence

Mice were anesthetized with ketamine/xylazine 80/10 mg/kg i.p. and transcardially perfused with ice-cold PBS followed by 4% paraformaldehyde (PFA; PanReac AppliChem ITW Reagents, Glenview, IL, USA) in PBS. Brains were carefully extracted and post-fixed overnight in 4% PFA at 4°C and then conserved in 30% sucrose (Merck KGaA, Darmstadt, Germany) for 1 week. Serial coronal brain slices (thickness: 40 μm) were cut with a freezing sliding microtome *HM 400* (MICROM International GmbH) and stored at-20 °C in cryoprotectant solution (30% glycerol and 30% ethylene glycol in 50 mM PBS).

Free-floating slices were washed three times with PBS. For c-Fos staining and antigen retrieval, slices were incubated in 10 mM sodium citrate (pH 8) for 30 minutes at 60°C. For Aβ immunostaining, slices were incubated in 70% formic acid for 10 minutes at room temperature (RT) to expose the Aβ epitope. All slices were then incubated in blocking solution (PBS containing 0.3% Triton X-100, 0.1% BSA, and 2% normal donkey serum) for 2 hours at RT. Following blocking, sections were incubated overnight at 4°C with the following primary antibodies: anti-Aβ clone 6E10 (1:200, cat# 803001, BioLegend), anti-NeuN (1:1000, cat# MAB377, Sigma-Aldrich), anti-c-Fos (1:1000 cat# 226308, Synaptic Systems), anti-IBA1 (1:1000, cat# 019-19741, Fujifilm Wako for double immunofluorescence with 6E10 and cat# HS-234308, Synaptic Systems, for double immunofluorescence with V-Glut1 and PSD-95 proteins), anti-V-Glut1 (1:1000, cat# MAB5502, Merck Millipore), anti-PSD95 (1:1000, cat# MA1-045, Thermo Fisher Scientific). Next day, slices were washed with PBS and incubated for 2 h at RT with the corresponding secondary antibodies Alexa Fluor donkey anti-mouse 488 (1:200, cat# A-21202), Alexa Fluor goat anti-guinea pig 488 (1:500, cat# A-11073), Alexa Fluor donkey anti-mouse 546 (1:500, cat#A-10036), Alexa Fluor goat anti-rabbit 568 (1:250, cat# A-11011), (all from Thermo Fisher Scientific). Finally, sections were washed with PBS and mounted with DAPI Fluoromount-G^®^ Mounting Medium (Southern Biotech, Birmingham, AL, USA).

To ensure consistent immunostaining, all sections were processed together under identical conditions. Images for amyloid plaques and c-Fos expression analysis were acquired using the Vectra Polaris scanner (Perkin Elmer). For microglial amyloid plaque coverage and synaptic material phagocytosis, images were captured using the LSM 800 Confocal Microscope (Leica Microsystems). Fluorescent signal analysis was performed using *ImageJ* software (version 1.48; NIH, Bethesda, MD, USA).

### 2.8. Quantification of Aβ1-42 levels in brain cortex

20 mg of brain cortex were homogenized in 8 volumes of cold 5 M guanidine-HCl in 50 mM Tris buffer. The homogenate was incubated for 3 h at RT on an orbital shaker and then, it was diluted ten-fold with cold PBS supplemented with 1X protease inhibitor cocktail (Calbiochem, Merck KGaA). Samples were centrifuged 20 min at 16,000 rcf at 4 °C and the supernatant was diluted 1:1000 for 5-month-old mice and 1:2000 for 8-month-old mice with Standard Diluent Buffer provided with the ELISA kit. Fifty microliters of the resultant solution were assayed using the *Ultrasensitive Amyloid-β 42 Human ELISA Kit* (cat# KHB3544; Invitrogen, Thermo Fisher Scientific) following the manufacturer’s instructions. The absorbance of each sample was read at 450 nm using the *Multiskan EX Microplate Reader* (Thermo Fisher Scientific). Each sample was analyzed in duplicate.

### 2.9. Statistical analysis

Normality was first checked by Shapiro-Wilk test. If the samples followed a normal distribution, the corresponding parametric statistical test was used; however, when the distribution was non-normal, the nonparametric test was performed. In the habituation and acquisition phases of the MWM, “AD and SIRT2 genotype” effects were analyzed by repeated-measures two-way ANOVA followed by multiple comparisons with Tukey’s test. In the case of LTP experiments, one-way ANOVA followed by multiple comparisons with Tukey’s test was performed. The rest of the behavioral tests and biochemical results were analyzed using two-way ANOVA followed by multiple comparisons with Tukey’s test. Post hoc test was applied only if F on interaction was significant. In figure legends, the F values represent the F of interaction followed by the p-value of the corresponding post hoc test. In those cases where the F of interaction was not statistically significant, the F value shown represents the main effect observed: AD or SIRT2 genotype. In those experiments that there were only two independent groups, the unpaired parametric Student’s t test or the nonparametric Mann–Whitney U test was performed. Survival curve was analyzed by Log-rank test. All results were expressed as mean ± standard error of the mean (SEM), and differences among groups were considered statistically significant at p < 0.05. All the statistics were performed by *GraphPad Prism* software (version 6.01; San Diego, CA, USA).

## 3. Results

### 3.1. Microglial deficiency of SIRT2 does not exert protective effects on AD pathology in 8-month-old APP/PS1 mice

To evaluate whether microglial SIRT2 deficiency could improve AD pathology, behavioral and molecular characterization was performed in 8-month-old APP/PS1 animals, as the most relevant symptoms of AD are already well established in this model.

Firstly, spontaneous motor activity was measured to exclude any influence of possible alterations in the basal motor activity of the animals in the subsequent behavioral tests. No differences were found in distance moved and velocity between the four groups (Figure 1A).

**Figure 1.**
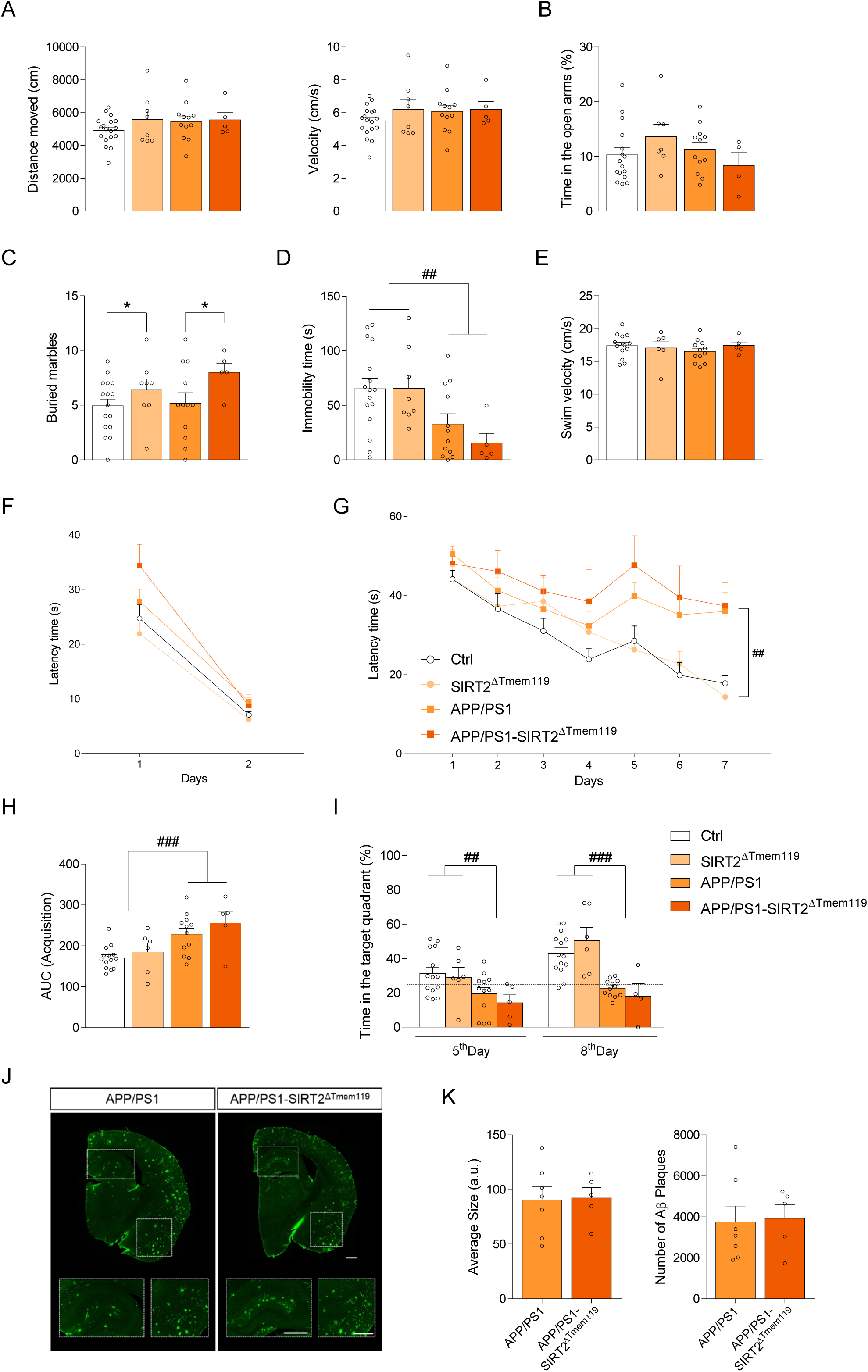
Microglial SIRT2 deficiency does not exert protective effects on AD pathology in 8-month-old APP/PS1 mice. **(A)** Distance moved (left) and velocity (right) of the spontaneous motor activity test. **(B)** Elevated plus maze, **(C)** marble burying test (F = 4.935, *p < 0.05, main effect of SIRT2 genotype, two-way ANOVA) and **(D)** forced swim test (F = 12.07, ##p < 0.01, main effect of AD, two-way ANOVA). Swim velocity **(E)** and escape latency **(F)** of the habituation phase of the MWM. Acquisition phase of the MWM: escape latency (F = 6.822, ##p < 0.01, Repeated Measures two-way ANOVA) **(G)** and area under the curve (AUC) of the acquisition curve (F = 15.34, ###p < 0.001, main effect of AD, two-way ANOVA) **(H)**. **(I)** Representation of the percentage of time spent in the correct quadrant in the retention phase of the MWM (5^th^ Day: F = 8.506, ##p < 0.01, main effect of AD; 8^th^ Day: F = 33.54, ###p < 0.01, main effect of AD, two-way ANOVA). Dotted line represents the 25% of time spent in the target quadrant (n = 5-18 animals per group). **(J)** Representative sections of Aβ plaques stained with 6E10 antibody (green) in brain slices of 8-month-old APP/PS1 and APP/PS1-SIRT2^ΔTmem119^ mice. Scale bar = 500 μm. **(K)** Amyloid burden quantification (n = 5-7 mice per group, 2 sections including hippocampus and cortex per animal): average size (left) and number of Aβ plaques (right). Results are shown as mean ± SEM. a.u.: arbitrary units.

Accumulating evidence suggests that microglia play a crucial role in sensing depression-related stressors and triggering immune responses that contribute to the development of depression (for a review, see [47]). To evaluate whether SIRT2 deficiency on microglia would affect anxiety and depressive-like states, EPM, Marble burying test and FST were performed. Regarding EPM, no significant differences were observed among all four groups (Figure 1B). Interestingly, SIRT2^ΔTmem119^ and APP/PS1-SIRT2^ΔTmem119^ mice buried more marbles compared to corresponding control and APP/PS1 animals (Figure 1C), suggesting an increase in anxiety-like behavior. Regarding FST, a main effect of AD genotype was observed, as APP/PS1 and APP/PS1-SIRT2^ΔTmem119^ showed a reduced immobility time compared to control animals, which could indicate an altered behavioral response in these groups that is not affected by microglial SIRT2 deficiency (Figure 1D).

Next, mice were subjected to the MWM task for learning and memory evaluation. On the habituation phase, all mice had the same swimming speed (Figure 1E), exhibited a normal swimming pattern and were able to reach the visible platform (Figure 1F). On the acquisition phase, the time spent to find the platform was significantly higher in APP/PS1 and APP/PS1-SIRT2^ΔTmem119^ mice (Figure 1G-H). On 5^th^ and 8^th^ day, when memory retention was evaluated, APP/PS1 and APP/PS1-SIRT2^ΔTmem119^ mice spent less time in the target quadrant than the corresponding control animals, indicating that these mice show cognitive deficiencies (Figure 1I). These results clearly demonstrate that microglial SIRT2 deficiency does not induce any improvement in the cognitive dysfunction of 8 month-old APP/PS1 mice.

Furthermore, histological analysis of hippocampal sections showed no significant differences in Aβ plaque burden, average plaque size, or the number of Aβ plaques between APP/PS1 and APP/PS1-SIRT2^ΔTmem119^ mice (Figure 1J-K). These findings suggest that, contrary to our initial hypothesis, microglial SIRT2 deficiency does not provide any beneficial effects in APP/PS1 mice with an AD established pathology.

### 3.2. Microglial SIRT2 deficiency reduces the survival and worsens learning decline in 5 month-old APP/PS1 mice

We then investigated whether SIRT2 deficiency might influence the onset of symptoms in the early stages of the disease, specifically in 5-month-old mice, by delaying their progression (Figure 2A). The survival of the animals was monitored for the first 25 weeks after birth (Figure 2B). APP/PS1 transgenic mouse model has short lifespan due to seizure activity and seizure-related sudden death [48]. As expected, survival of APP/PS1 animals decreased after 10 weeks of age. Surprisingly, at 25 weeks of age, the mortality of APP/PS1-SIRT2^ΔTmem119^ was increased compared to APP/PS1 mice (48.3% vs 63% of survival rate, respectively), indicating that SIRT2 deficiency reduced the survival of APP/PS1 mouse model.

**Figure 2.**
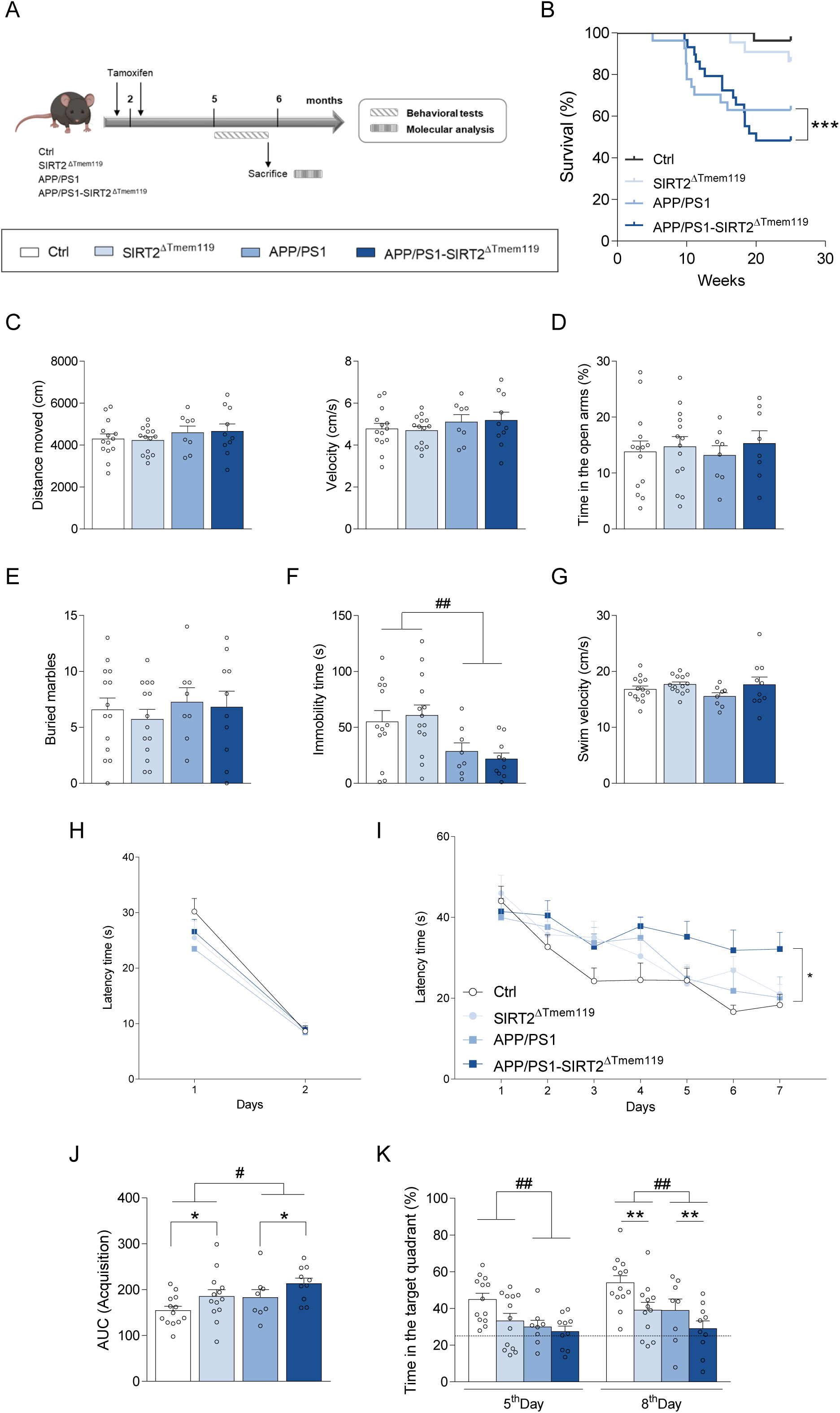
Microglial SIRT2 deficiency reduces the survival and worsens memory of 5 month-old APP/PS1 mice. **(A)** To achieve the cre-dependent deletion of *Sirt2* gene, tamoxifen was administered to male and female mice when they were 6 and 10 weeks old. At the age of 5 months, mice were subjected to a series of behavioral tests prior to postmortem molecular analysis where amyloid-β pathology and synaptic function were evaluated. **(B)** Survival analysis of APP/PS1-SIRT2^ΔTmem119^ mouse model during the first 25 weeks of age (***p < 0.001, Log-rank test) (Ctrl, n = 27; SIRT2^ΔTmem119^, n = 22; APP/PS1, n = 27; APP/PS1-SIRT2^ΔTmem119^, n = 29). **(C)** Distance moved (left) and velocity (right) of the spontaneous motor activity test. **(D)** Elevated plus maze, **(E)** marble burying test and **(F)** forced swim test (F = 12.49, ##p < 0.01, main effect of AD, two-way ANOVA). Swim velocity **(G)** and escape latency **(H)** of the habituation phase of the MWM. Acquisition phase of the MWM: escape latency (F = 3.555, *p < 0.05, Repeated Measures two-way ANOVA) **(I)** and area under the curve (AUC) of the acquisition curve (F = 5.142, *p < 0.05, main effect of SIRT2 genotype; F = 4.433, #p < 0.05 main effect of AD, two-way ANOVA) **(J)**. **(K)** Representation of the percentage of time spent in the correct quadrant in the retention phase of the MWM (5^th^ Day: F = 7.915 ##p < 0.01, main effect of AD; 8^th^ Day: F = 7.440, **p < 0.01, main effect of SIRT2 genotype; F = 7.648, ##p < 0.01, main effect of AD, two-way ANOVA). Dotted line represents the 25% of time spent in the target quadrant (n = 8-14 animals per group). Results are shown as mean ± SEM.

We next assessed spontaneous motor activity and found no differences in distance moved or velocity among the four groups (Figure 2C), ruling out any influence of alterations in basal motor activity. Regarding anxiety-like behavior, no differences were observed in the time spent in the open arms during the EPM test (Figure 2D) and the number of buried marbles was similar across all groups (Figure 2E). Additionally, consistent with findings in older animals, APP/PS1 mice exhibited reduced immobility time in the FST compared to control animals, indicating a depressive-like phenotype, with no observed effects of microglial SIRT2 deficiency (Figure 2F). These results suggest that microglial SIRT2 is not involved in regulating depression-or anxiety-related behaviors in 5 month-old mice.

Mice were then subjected to the MWM task to assess the impact of microglial SIRT2 deficiency on spatial learning and memory. During the habituation phase, no differences in swimming speed were observed (Figure 2G) and all mice successfully reached the visible platform (Figure 2H). During the acquisition phase, the time to find the hidden platform decreased significantly each day in the control, SIRT2^ΔTmem119^ and APP/PS1 mice. However, APP/PS1-SIRT2^ΔTmem119^ mice exhibited a higher escape latency, indicating learning and memory deficits at this age (Figure 2I). Notably, when the area under the acquisition curve (AUC) was analyzed, a significant main effect of the SIRT2 genotype was detected (Figure 2J), suggesting that microglial SIRT2 deficiency impaired learning capacity.

As shown in Figure 2K, during the first probe trial (5^th^ day), APP/PS1 mice spent less time in the target quadrant, indicating cognitive deficiencies. No significant differences were observed between control and SIRT2^ΔTmem119^ groups. However, consistent with the acquisition phase results, the probe trial on the 8^th^ day revealed that APP/PS1-SIRT2^ΔTmem119^ mice still failed to remember the location of the platform, while the APP/PS1 group, with nearly 40% of time spent in the target quadrant, did (Figure 2K). Interestingly, SIRT2^ΔTmem119^ mice also showed worse memory retention compared to control mice during this phase (Figure 2K).

### 3.3. Microglial SIRT2 deficiency accelerates Aβ deposits in 5-month-old APP/PS1 mice

As shown in Figure 3A, 6E10 immunostaining was significantly increased in APP/PS1-SIRT2^ΔTmem119^, indicating a worsening of amyloid pathology. Further analysis of stained amyloid plaques revealed that, while there were no differences in the total plaque count in the analyzed histological sections (Figure 3B), the average plaque size was significantly larger in APP/PS1-SIRT2^ΔTmem119^ animals (Figure 3C). Additionally, elevated levels of Aβ1-42, measured by ELISA, were also detected in microglial SIRT2 deficient APP/PS1 mice (Figure 3D). No Aβ was detected in control and SIRT2^ΔTmem119^ littermates (data not shown). The accelerated amyloid pathology in APP/PS1-SIRT2^ΔTmem119^ brains was accompanied by a significant increase in the gene expression of the pro-inflammatory cytokines *Il-1β* (Figure 3E) and *Il-6* (Figure 3F), with no significant differences in *Tnf-α* expression (Figure 3G).

**Figure 3.**
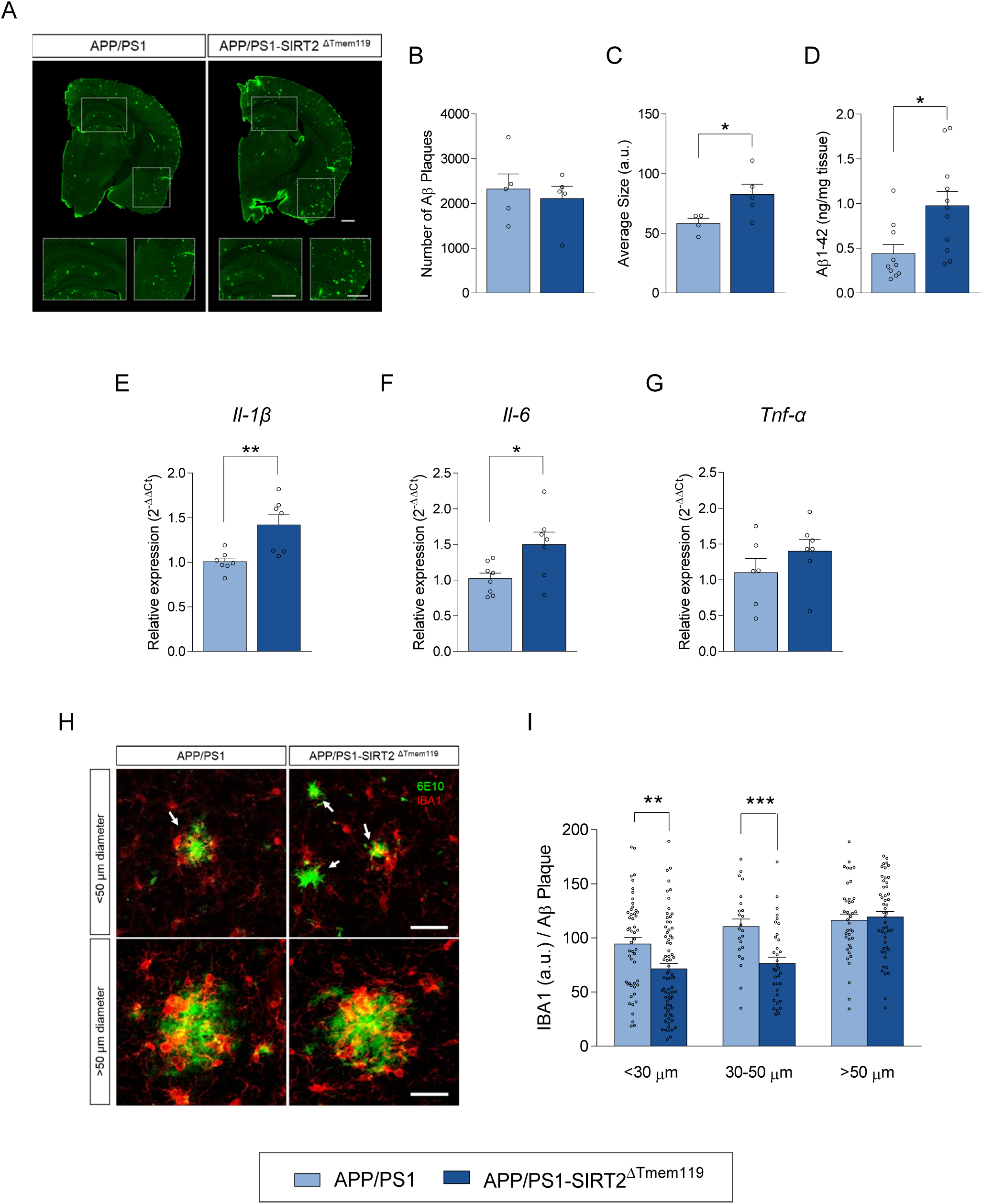
Microglial SIRT2 deficiency exacerbates amyloid pathology in 5 month-old APP/PS1 mice. **(A)** Representative sections of Aβ plaques stained with 6E10 antibody (green) in brain slices of 5-month-old APP/PS1 and APP/PS1-SIRT2^ΔTmem119^ mice. Scale bar = 500 µm. Amyloid burden quantification (n = 5 mice per group, 2 sections including hippocampus and cortex per animal): number **(B)** and average size of Aβ plaques (*p < 0.05, Student’s t test) **(C)**. **(D)** Levels of Aβ1-42 in the cortex of APP/PS1 and APP/PS1-SIRT2^ΔTmem119^ mice measured by ELISA (*p < 0.05, Student’s t test) (n = 11 mice per group). Gene expression of *Il-1β* (**p < 0.01, Student’s t test) **(E)**, *Il-6* (*p < 0.05, Student’s t test) **(F)** and *Tnf-α* **(G)**. *Gapdh* was used as internal control (n = 7-8 mice per group). **(H)** Representative sections of Aβ plaques stained with 6E10 antibody (green) and microglial cells stained with IBA1 antibody (red) in brain slices of 5-month-old APP/PS1 and APP/PS1-SIRT2^ΔTmem119^ mice. Arrows indicate Aβ plaques smaller than 50 µm of diameter. Scale bar = 30 µm. **(I)** IBA1 expression surrounding Aβ plaques. Aβ plaques were divided by size in three groups: 0-30 µm, 30-50 µm and bigger than 50 µm of diameter (**p < 0.01, ***p < 0.001, Student’s t test) (n = 50-60 plaques per group). Results are shown as mean ± SEM. a.u.: arbitrary units.

As Aβ clearance is mediated by microglia, we next analyzed the coverage of microglia to Aβ plaques. In APP/PS1 mice, a marked microglia “clustering” phenotype was observed since microglia were found to surround Aβ plaques (Figure 3H). Noteworthy, in APP/PS1-SIRT2^ΔTmem119^ mice the percentage of Aβ plaque-coverage by microglia was decreased in plaques smaller than 50 μm of diameter (Figure 3I). This reduction in microglial coverage suggests that SIRT2 deletion may lead to microglial dysfunction, impairing their ability to recognize these incipient protein aggregates. Such dysfunction could, in turn, explain the increased size of amyloid plaques observed in APP/PS1-SIRT2^ΔTmem119^.

### 3.4. Transcriptomic analysis of SIRT2 deficient microglia revealed changes in aging and neuronal activity related genes

To investigate the impact of SIRT2 reduction on microglial gene expression and its potential role in exacerbating the AD phenotype, we performed bulk RNA sequencing on microglial samples isolated from the hippocampus of 5 month-old control, SIRT2^ΔTmem119^, APP/PS1 and APP/PS1-SIRT2^ΔTmem119^ mice (Figure 4A).

**Figure 4.**
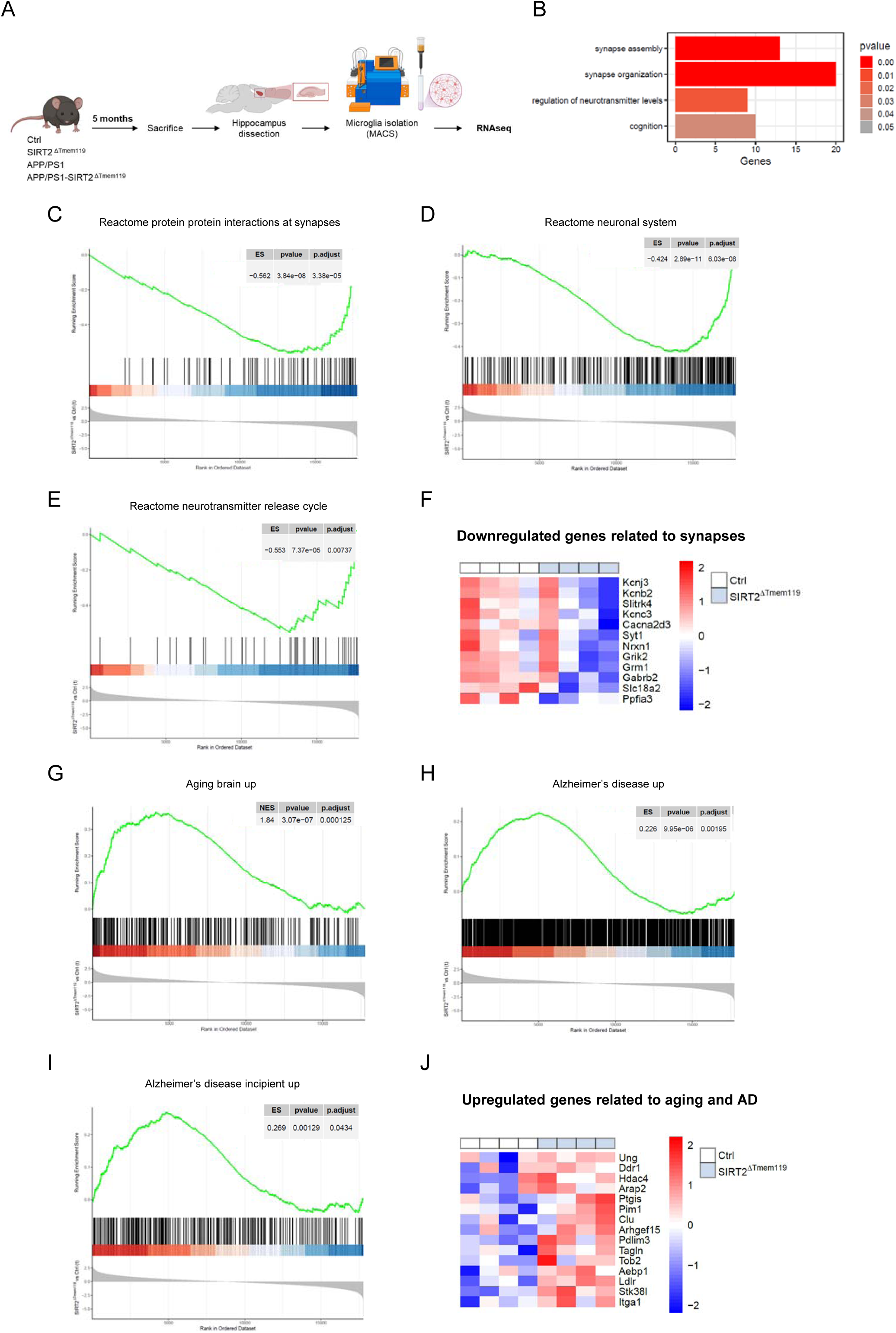
SIRT2 deficient microglia exhibit alterations in the expression of genes associated with aging and neuronal activity. **(A)** Five-month-old mice were sacrificed and microglia from the hippocampus were isolated to perform bulk RNA sequencing. **(B)** GO Biological Processes with the highest significant fold enrichment when control and SIRT2^ΔTmem119^ groups were compared. **(C-E)** Gene Set Enrichment Analysis (GSEA) indicating a repression of genes related to synaptic interaction **(C)**, neuronal system **(D)** and neurotransmitter release **(E)** in SIRT2^ΔTmem119^ microglia. **(F)** Heatmap showing the most downregulated genes associated with synapses in SIRT2^ΔTmem119^ microglia. **(G-I)** Gene Set Enrichment Analysis (GSEA) indicating an upregulation of genes related to aging **(G)** and Alzheimer’s disease **(H-I)** in SIRT2^ΔTmem119^ microglia. **(J)** Heatmap showing the most upregulated genes associated with aging and Alzheimer’s disease (AD) in SIRT2^ΔTmem119^ microglia.

Initially, we compared the control and SIRT2^ΔTmem119^ groups to determine how the deletion of SIRT2 affects the transcriptional profile of microglia in physiological conditions (Figure 4B-J). The results showed that a total of 526 genes (200 up-regulated genes and 326 down-regulated genes) were differentially expressed in SIRT2^ΔTmem119^ mice compared to the control group. Interestingly, gene ontology (GO) analysis revealed that the GO Biological Processes with the highest significant fold enrichment were related to synapse assembly, regulation and organization (Figure 4B). Furthermore, Gene Set Enrichment Analysis (GSEA) indicated a repression of genes related to synaptic interaction, neuronal system and neurotransmitter release (Figure 4C-E). Among the downregulated genes (Figure 4F) we found changes in *Kcnj3*, *Kcnb2*, *Kcnc3*, and *Cacna2d3,* involved in the modulation of ion flow across membranes; *Syt1*, *Nrxn1*, *Grik2*, *Grm1* and *Gabrb2* crucial for vesicle release and neurotransmitter signaling; and *Ppfia3* involved in cell signaling, interactions with extracellular matrix and adhesion processes.

Additionally, GSEA analysis also revealed an enrichment of aging and AD-associated genes in SIRT2^ΔTmem119^ microglia (Figure 4G-I). The upregulated genes (Figure 4J) include genes involved in inflammation and immune response (*Clu* and *Ptgis*), extracellular matrix interactions (*Ddr1* and *Itga1*), cytoskeletal dynamics, migration and cellular adhesion (*Arhgef15*, *Pdlim3*, *Tm4sf1* and *Stk38l*) and genes linked to lipid homeostasis and cholesterol metabolism, which are crucial in neurodegenerative processes such as *Ldlr, Abcg1,* and *Cyp1b1*. These findings provide insights into the molecular alterations in SIRT2 deficient microglia that may contribute to age-related neurodegenerative changes.

Next, we further examined the impact of SIRT2 deficiency in microglia in the response to changes occurring in the pathological context of the APP/PS1 model (Figure 5). Pairwise analyses revealed more differentially expressed genes between APP/PS1- SIRT2^ΔTmem119^ and SIRT2^ΔTmem119^ (652 genes changed) than between Control and APP/PS1 groups (251 genes changed), suggesting a higher impact of AD pathology in a context of SIRT2 deficiency at this age (Figure 5A). When we further analyzed the functionality of these genes we found some functions like chromatin remodeling, innate immune response or cellular response to ketone or nitrogen changed in both conditions with AD (APP/PS1-SIRT2^ΔTmem119^ and APP/PS1) when compared with their corresponding non-AD control groups (SIRT2^ΔTmem119^ and Ctrl respectively) (Figure 5B). Noteworthy, in the context of SIRT2 deficiency, we found many cellular functions affected in the presence of the APP/PS1 mutations that are not altered when SIRT2 expression is not modulated. Some of these functions include cellular response to stress, Wnt signaling pathway, sterol transport, regulation of stress-activated MAPK cascade, response to endoplasmic reticulum stress, VEGFA VEGFR2 signaling, SNARE interactions in vesicular transport and vesicle-mediated transport. Some other functions, however, exhibit changes in the APP/PS1 model when compared to the control group, but these alterations are not observed in the comparison between both groups in the absence of SIRT2. This suggests that SIRT2 deficiency may prevent the activation or repression of certain functions that are normally modified in the context of early AD or that these functions were already altered at baseline in the SIRT2^ΔTmem119^ group (i.e. cytoskeletal regulation or lipid metabolism).

**Figure 5.**
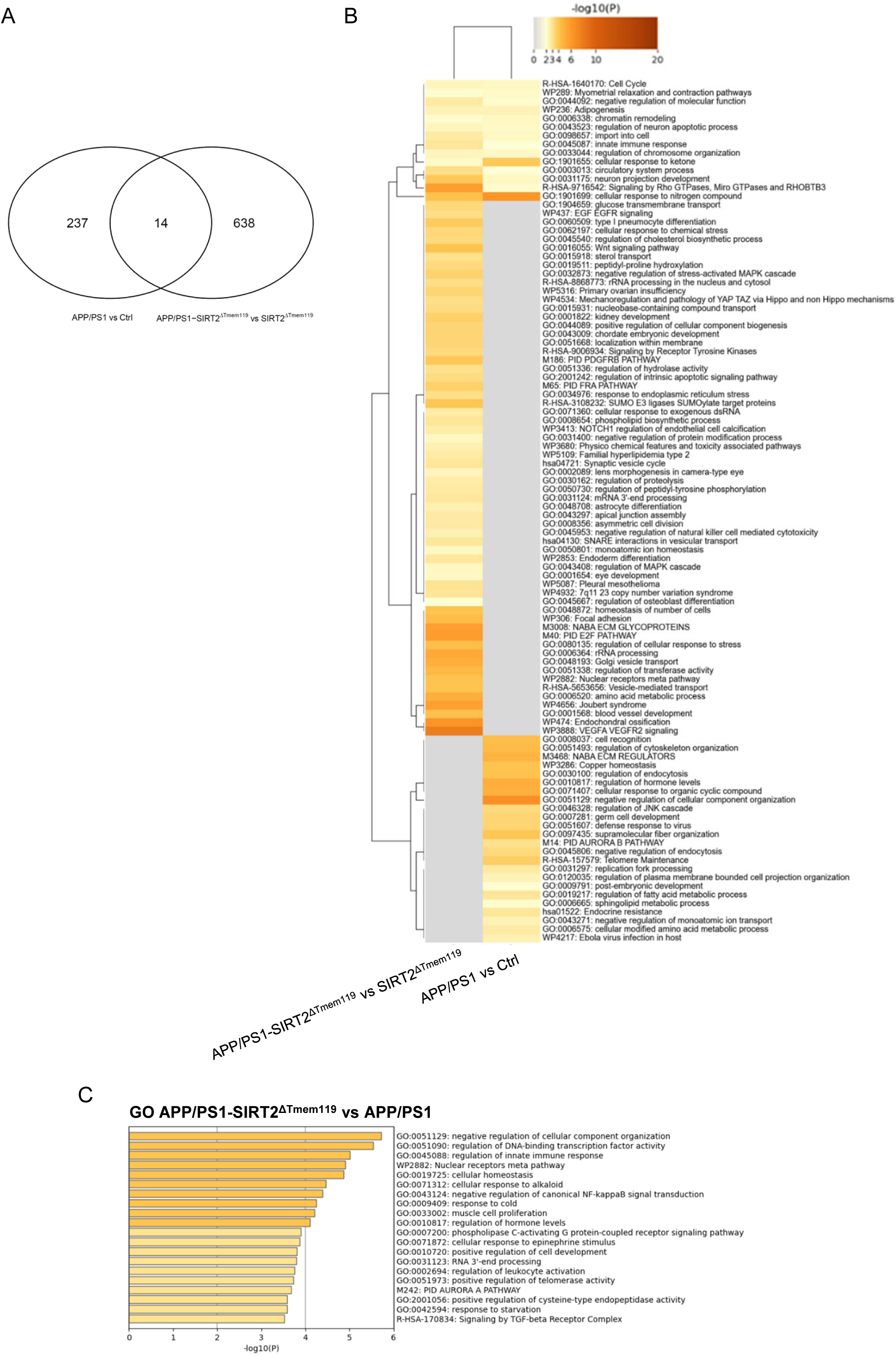
Microglial deficiency of SIRT2 modifies the transcriptome in the context of Alzheimer’s disease pathology. **(A)** Venn diagram illustrating the comparison of differentially expressed genes between APP/PS1 versus Control microglia and APP/PS1 versus APP/PS1-SIRT2^ΔTmem119^ microglia. Overlapping areas represent shared genes between the two comparisons, while non-overlapping regions indicate genes specific to each group. **(B)** Heatmap of the Gene Ontology (GO) functional analysis of differentially expressed genes in the comparisons APP/PS1 versus Control microglia and APP/PS1 versus APP/PS1-SIRT2^ΔTmem119^ microglia. Color intensity represents the statistical significance (-log10 p-value) of each enriched GO term. **(C)** Top 20 GO Biological Processes altered in the comparison between APP/PS1-SIRT2^ΔTmem119^ and APP/PS1 groups.

When further analyzing the cellular functions associated with the differentially expressed genes in the APP/PS1- SIRT2^ΔTmem119^ compared to the APP/PS1 group, we observed that the top 20 functions, as depicted in Figure 5C, encompass a range of critical biological processes. These include the negative regulation of cellular component organization, modulation of DNA-binding transcription factor activity, regulation of innate immune response, cellular homeostasis, negative regulation of canonical NF-kB signal transduction and signaling by the TGF-beta receptor complex. These findings underscore the impact of SIRT2 deficiency on cellular functions that are essential for maintaining cellular homeostasis and response to stress in the context of AD, highlighting potential pathways that may contribute to neurodegenerative processes.

### 3.5. Microglial SIRT2 deficiency impairs neuronal activity and LTP

To validate the transcriptomic results related to neuronal activity and synaptic transmission, we next assessed histologically the neuronal consequences of microglial SIRT2 deficiency. We first quantified c-Fos immunoreactivity in the hippocampus and observed a significant reduction in c-Fos+ cells in both SIRT2^ΔTmem119^ and APP/PS1- SIRT2^ΔTmem119^ groups compared to control and APP/PS1 animals respectively suggesting a decrease in neuronal activity (Figure 6A). As shown in Figure 6B, no significant differences were found in the immunofluorescence intensity of the neuronal marker NeuN across all four groups, suggesting that neurotoxicity is not the mechanism driving the detrimental effects of microglial SIRT2 deletion. We next investigated the morphology of the dendritic spine in our experimental conditions. Interestingly, while the number of dendritic spines remained unchanged (Figure 6C), Golgi-Cox-stained hippocampal slices revealed alterations in the morphology of the dendritic spines (Figure 6D-F). Specifically, APP/PS1-SIRT2^ΔTmem119^ mice exhibited a reduction in thin spine density (Figure 6D) and an increase in mushroom-type spines (Figure 6E). It is worthy to mention that, while a moderate increase in mushroom spines may be associated with improved LTP and cognitive function, an excessive or dysregulated increase can have detrimental effects on synaptic plasticity and neuronal function [49,50]. Thus, we next investigated the impact of microglial SIRT2 deletion on synaptic plasticity by assessing LTP in *ex vivo* hippocampal slices. Supporting previous studies in whole body SIRT2 knockout mice [51] and in myeloid specific SIRT2 deficient mice under inflammatory conditions [52], and consistent with the cognitive impairments observed in the MWM, LTP in SIRT2^ΔTmem119^ and APP/PS1-SIRT2^ΔTmem119^ mice was substantially reduced compared to control mice (Figure 6G). Notably, at this age, LTP in APP/PS1 slices is not yet fully impaired. These findings suggest that microglial SIRT2 deletion has a pronounced effect on synaptic plasticity, potentially accelerating the onset of the pathology in the APP/PS1 mouse model.

**Figure 6.**
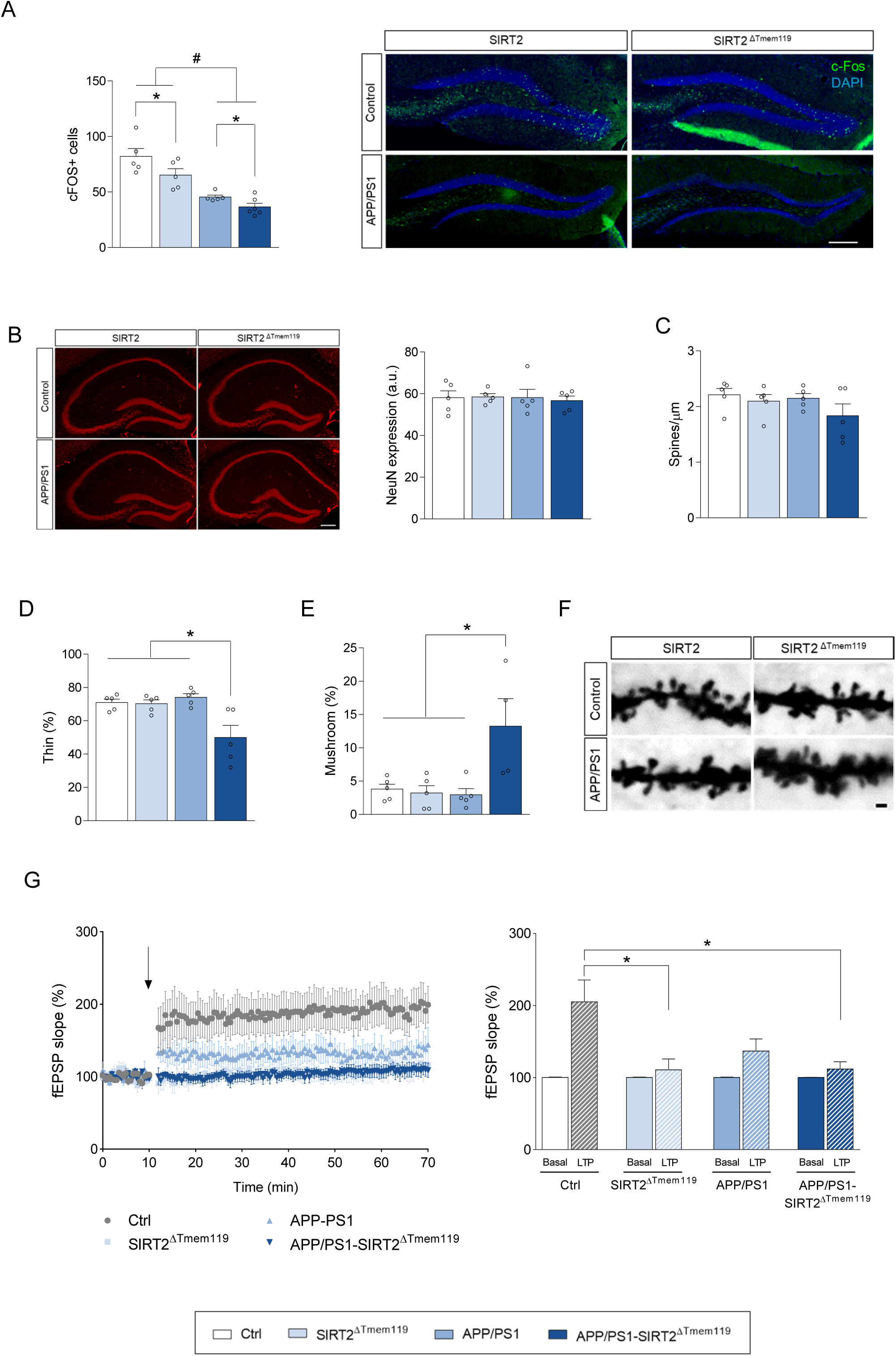
Microglial deficiency of SIRT2 impairs neuronal activity and alters spine morphology. **(A)** Quantification of c-Fos positive cells (left) and representative c-Fos staining (green) in the dentate gyrus of the hippocampus (right) (n = 6 mice per group; 1 section per animal). Scale bar = 300 µm. **(B)** Representative hippocampal sections of NeuN staining (red; left) and quantitative measurement of NeuN expression in the hippocampus (right) (n = 5 mice per group; 3 sections per animal including CA1, CA3 and DG). Scale bar = 200 µm. **(C)** Spine density. **(D)** Percentage of thin (F = 4.819, *p < 0.05, two-way ANOVA followed by Tukey’s test) and **(E)** mushroom type spines (F = 6.765, *p < 0.05, two-way ANOVA followed by Tukey’s test) (n = 5 animals per group). **(F)** Representative images of Golgi-Cox-stained dendritic spines Scale bar = 1 µm. **(G)** (Left) Time course of mean fEPSP slope in hippocampal slices from WT, SIRT2^ΔTmem119^, APP/PS1 and APP/PS1-SIRT2^ΔTmem119^ mice in basal conditions and following induction of theta burst stimulation at 10 minutes (arrow). (Right) Average relative changes of fEPSP slopes in hippocampal mice slices before (basal) and after 60 minutes (LTP) application of theta burst (F = 4.512, *p < 0.05, one-way ANOVA followed by Tukey’s test) (n = 3-5 animals per group, 3-4 slices per animal). Results are shown as mean ± SEM. a.u.: arbitrary units.

In order to reinforce these studies we analyzed microglial phagocytosis of synaptic proteins using a double immunostaining for IBA1 (microglial marker) and either VGLUT1 (presynaptic terminals) or PSD95 (postsynaptic elements). Confocal microscopy analysis of internalized synaptic material within IBA1+ cells in the hippocampus revealed no significant differences in VGLUT1 phagocytosis by microglia (Figure S2). However, SIRT2^ΔTmem119^ microglia exhibited a significant increase in PSD95 colocalization compared to control microglia (Figure 7A-C), supporting the hypothesis that microglial SIRT2 deficiency impacts on microglia-neuron interactions at synaptic level. In contrast, this increased engulfment was not observed in APP/PS1-SIRT2^ΔTmem119^ animals, likely due to the fact that a significant portion of microglia are clustered around large amyloid plaques in these animals (as shown in Figure 3H). This shift suggests an alteration in microglial functionality, with their activity directed more toward plaque-associated tasks than synaptic modulation in this pathological context.

**Figure 7.**
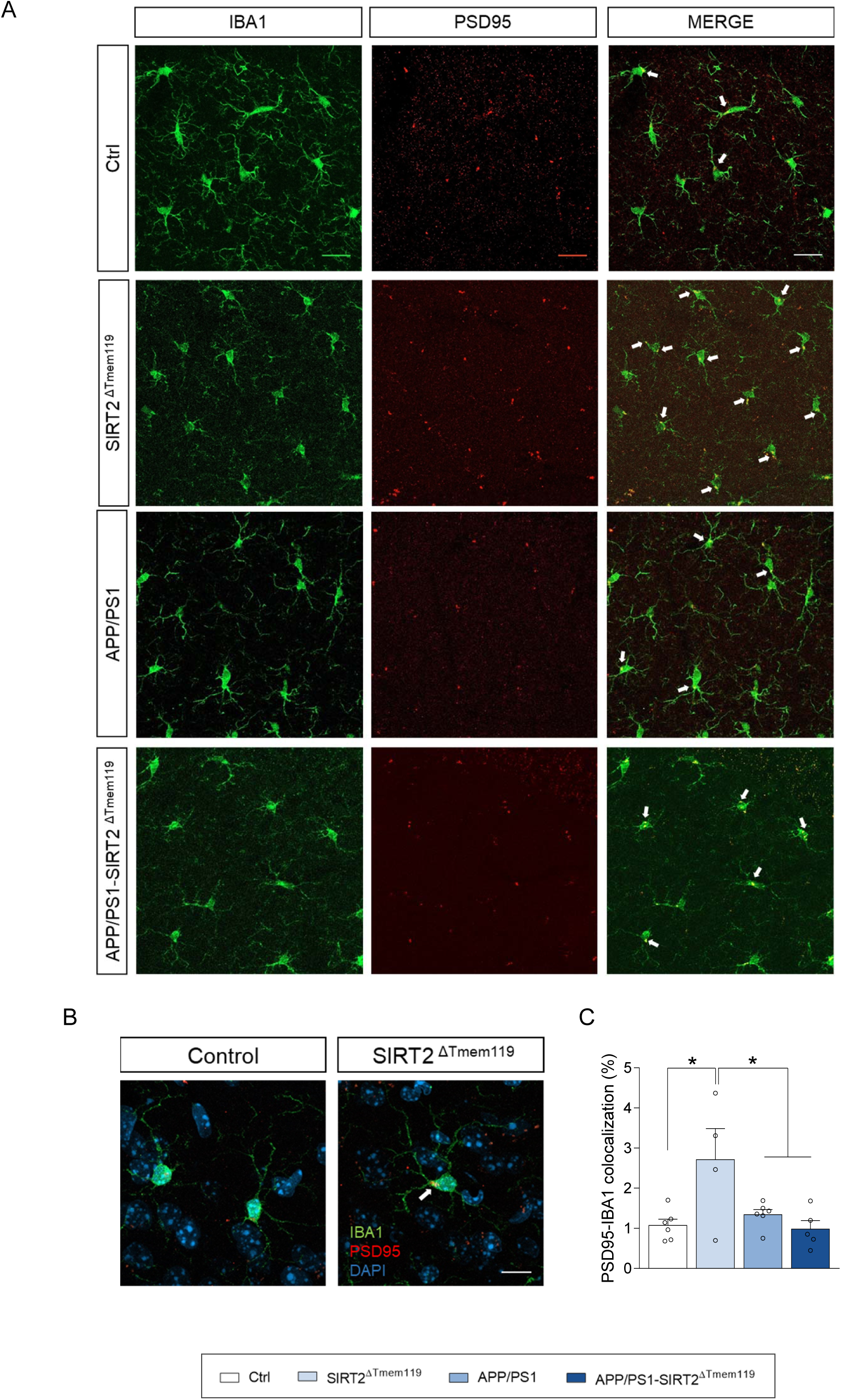
SIRT2 deficiency enhances microglial phagocytosis of PSD95. **(A)** Representative images of microglial cells stained with IBA1 antibody (green, microglial marker) and PSD95 (red, postsynaptic marker). Arrows indicate areas of colocalization between IBA1 and PSD95. Scale bar = 20 µm. **(B)** Higher magnification of representative microglia from control and SIRT2^ΔTmem119^ mice, showing increased of PSD95 engulfment in SIRT2 deficient microglia (arrow). Scale bar = 10 µm. **(C)** Quantitative measurement of IBA1-PSD95 colocalization in brain slices from 5-month-old mice (n = 4-6 mice per group; 1 section per animal). Results are shown as mean ± SEM.

## 4. Discussion

SIRT2 has been identified as a key regulator of aging [3,6] and its inhibition has shown beneficial effects in age-related neurodegenerative diseases such as AD [13–16,53]. However, the potential adverse effects associated to its peripheral pharmacological inhibition [14] necessitate careful consideration to ensure the development of safe and effective therapeutic strategies. In this study, we propose that targeted inhibition of SIRT2 within the CNS, specifically in microglia, may offer a safer and more effective approach for the treatment of AD.

However, our findings demonstrate that microglial SIRT2 deficiency does not confer protective effects against AD pathology in APP/PS1 mice and even exacerbates certain aspects of the disease in its early stages. Transcriptomic analyses revealed that SIRT2-deficient microglia express gene profiles associated with aging and AD, along with alterations in neuronal activity evidenced by a reduction in c-Fos+ cells in the hippocampus. Correspondingly, these mice showed worse cognitive performance, increased microglial engulfment of postsynaptic elements and impaired LTP in the hippocampus. In a pathological context, microglial SIRT2-deficient APP/PS1 mice exhibited reduced survival rates, accelerated cognitive decline, increased amyloid plaque deposition, and elevated levels of pro-inflammatory cytokines at early disease stages. These changes were accompanied by impaired synaptic function, alterations in dendritic spines morphology and disrupted LTP in the hippocampus.

These findings contrast with those previous studies showing that SIRT2 pharmacological inhibition or whole-body genetic deletion enhances learning and memory and reduces mortality and amyloid pathology in mouse models of AD [13–16]. Bai et al. [13] and Wang et al. [16] demonstrated that whole-body SIRT2 knockout or inhibition reduces amyloid pathology by shifting APP processing toward the non-amyloidogenic pathway, either through increased APP acetylation or inhibition of BACE1 activity. In contrast, a recent study aligns with our findings, showing that SIRT2 deficiency can exacerbate amyloid pathology [54]. Chen et al. have demonstrated that SIRT2 inhibition or knockdown reduces ApoE secretion in the BV2 microglial cell line, which in turn diminishes the degradation of extracellular Aβ levels [54]. ApoE, secreted by microglia, interacts with Aβ in the extracellular space of the brain, facilitating Aβ degradation and clearance [55]. Notably, we have also observed a reduction in plaque-associated microglia, specifically around small and medium plaques. In this sense, recent studies highlight that microglial containment and encapsulation of Aβ plaques are crucial microglial functions in response to Aβ mediated pathology [56–60]. Therefore, we propose that neuron SIRT2 deletion may be beneficial by reducing Aβ production [13,16], while microglia-specific SIRT2 deletion may impair Aβ degradation ([54]; present study), thereby exacerbating the pathology. This discrepancy underscores the need for conditional and cell-specific transgenic models to accurately assess the role of SIRT2 in aging and disease.

Another intriguing finding from the present study is that SIRT2 deficiency in microglia is associated with an increase in aging-related genes. While SIRT2 is often linked to aging and longevity, its role in promoting or preventing aging remains a topic of debate (for a review, see [3]). Our results corroborate previous studies that underscore the critical role of SIRT2 and the necessity of maintaining its expression across various cell types (such as macrophages, cardiomyocytes, and vascular smooth muscle cells) to mitigate age-related inflammation, insulin resistance, cardiac dysfunction, and vascular remodeling [20,22,23]. Notably, associations of premature senescence with microglia have been observed in the APP/PS1 mouse model and in postmortem AD samples [61,62]. Furthermore, evidence suggests that preventing microglial senescence can lead to reduced pathology, thereby highlighting the link between microglial activity and the early stages of AD [61]. Indeed, it has been proposed that microglia should be a primary focus for senolytic treatments in AD [62]. This therapeutic potential underscores the importance of further investigating the mechanisms underlying premature aging in microglia.

Our results also highlight the impact of microglia on synaptic plasticity, aligning with findings from numerous other studies [63]. We found that LTP recorded from hippocampal slices of microglial SIRT2-deficient mice was significantly impaired. This observation in microglial SIRT2-deficient APP/PS1 mice may be interpreted as a further indication of accelerated or aggravated AD pathology, as APP/PS1 mice of this age already exhibit some LTP impairment, which eventually progresses to complete loss as amyloid pathology and neuroinflammation advance. However, the mechanisms underlying LTP impairment in SIRT2^ΔTmem119^ brains remain unclear and warrant further investigation. Our findings align with previous studies that observed LTP impairments in whole body SIRT2 knockout mice [51] and in myeloid specific SIRT2 deficient mice under inflammatory conditions [52]. Wang and coworkers proposed that LTP alterations in SIRT2 knock out mice could result from enhanced acetylation and surface expression of AMPA receptors in neurons [51]; however, since in our model SIRT2 expression is unaffected in neurons, we can likely exclude this as a direct mechanism involved.

Microglia, considered the fourth component of the ‘quad-partite synapse’, influences neuronal communication by interacting with both presynaptic and postsynaptic elements [64]. Notably, we observed increased engulfment of the postsynaptic protein PSD95 in the hippocampus of SIRT2^ΔTmem119^ mice, which could provide a plausible mechanism underlying the synaptic impairments in these animals. Nonetheless, other mechanisms should also be explored. Potential candidates include molecules released by microglia under both physiological and pathological conditions such as TNF-α, IL-1β, IL-6, glutamate and reactive oxygen and nitrogen species, all of which have been shown to modulate synaptic transmission and plasticity [65–68]. In line with this, Sa de Almeida et al. rescued LTP impairment in myeloid-specific SIRT2 deficient mice using memantine, an NMDA antagonism [52], suggesting that SIRT2 deficiency may enhance NMDA-induced toxicity and impair LTP and cognition. Another plausible mechanism underlying LTP and synaptic alterations in our model could involve microglia-derived exosomes. In the CNS, exosomes play a key role in intercellular communication between glial cells and neurons, modulating neuroplasticity [69,70]. Interestingly, recent findings show that oligodendrocyte-derived exosomes contain high levels of SIRT2, and that its transfer from oligodendrocytes to neurons is critical for synaptic activity and neuroplasticity [71,72]. Given that microglia-derived exosomes mediate neuronal survival, neurite outgrowth, synaptic function and neuroinflammatory responses through enzymes, chaperones, membrane receptors, and miRNAs [73,74], we speculate that altered exosomal communication could underlie the LTP deficits observed in our model. Further studies are needed to characterize SIRT2 deficient microglia-derived exosomes to test this hypothesis.

In summary, we have demonstrated that SIRT2 plays a key role in microglial function and that its genetic deletion is associated with exacerbation of early AD pathology. Given that microglia are central to the etiology of numerous neurodegenerative diseases, our findings have significant implications: while previous studies have suggested a neuroprotective role for SIRT2 inhibition, our results indicate that the manipulation of SIRT2 expression and activity should be approached with caution. Further investigations into the specific functions of SIRT2 in different cell types will be necessary to establish a rational framework for its use as a therapeutic target.

## 6. Acknowledgements

The authors are grateful to Sandra Lizaso for her excellent technical assistance. We would also like to thank “Amigos de la Universidad de Navarra” and the Spanish Ministry of Universities for a fellowship to N.S-S and M.G-B.

The graphical abstract was created using biorender.com.

## 7. Author Contributions

N.S-S: methology, investigation, formal analysis, writing-original draft preparation, writing-review and editing. M.G-B: methology, investigation, formal analysis, writing-original draft preparation, writing-review and editing. M.A: methology, investigation. MC.M-D: methology, investigation, formal analysis. L.G-C: investigation, formal analysis. S.E: investigation, formal analysis. E.A-C: data curation, formal analysis. J.F-I: data curation, formal analysis. M.S: conceptualization, writing-review and editing. RM.T: conceptualization, writing-review and editing. E.S: data curation, formal analysis, writing-review and editing. ED.M: investigation, formal analysis, writing-review and editing. E.P: conceptualization, investigation, supervision, formal analysis, writing-original draft preparation, writing-review and editing, funding acquisition. All authors have reviewed and approved the final version of the manuscript for publication.

---

**Supplementary Figure 1.**
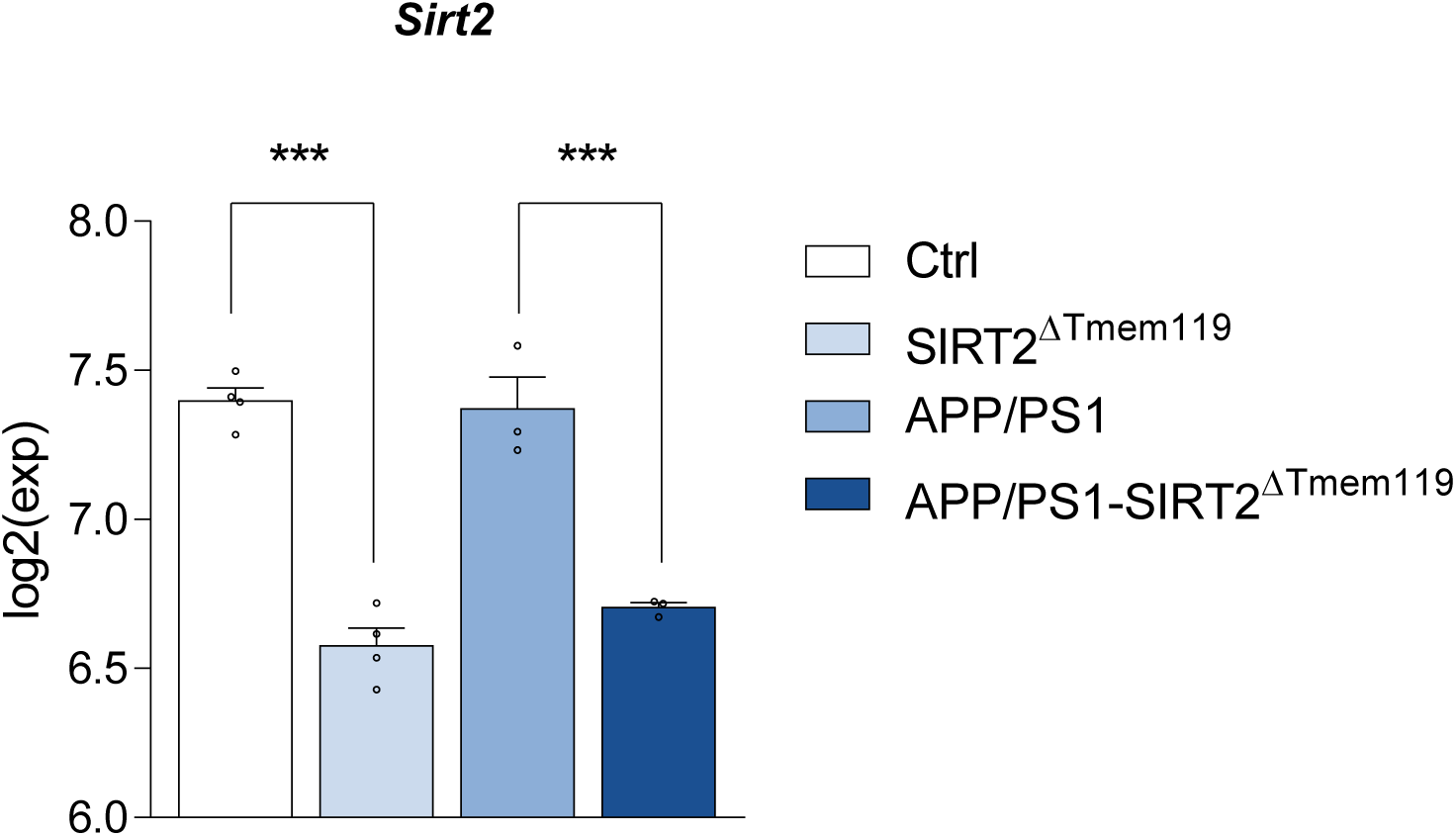
Sirtuin 2 expression in isolated microglia. Gene expression of *Sirt2* analyzed by bulk RNA sequencing in microglia isolated from the hippocampus of five-month-old mice.

**Supplementary Figure 2.**
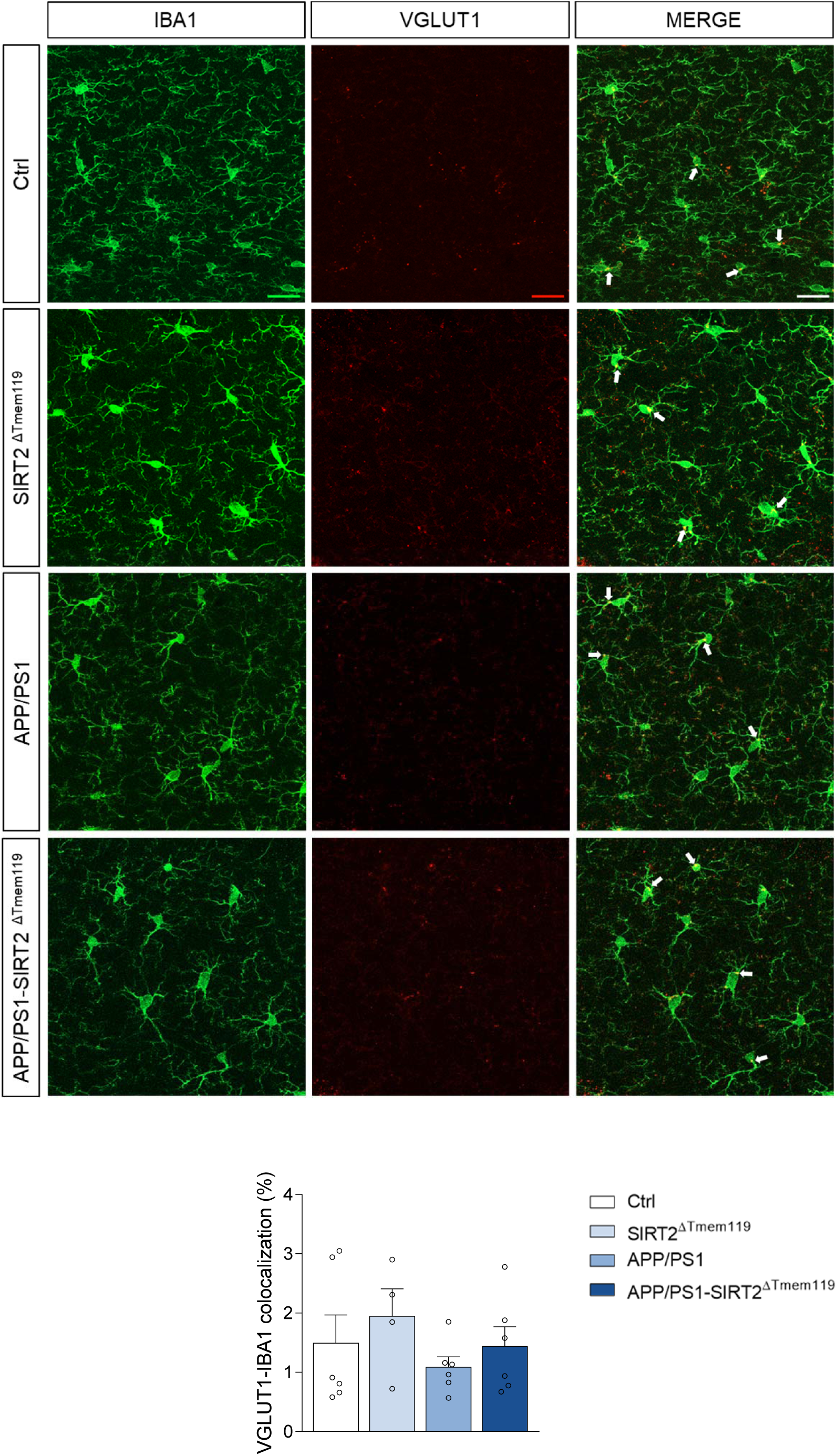
Microglial phagocytosis of presynaptic proteins (VGLUT1) Representative sections of microglial cells stained with IBA1 antibody (green) and VGLUT presynaptic marker (red) (top) and quantitative measurement of the colocalization of both markers in brain slices of 5-month-old mice (bottom) (n = 4-6 mice per group; 1 section per animal). Arrows indicate colocalization of IBA1 and VGLUT. Scale bar = 20 µm. Results are shown as mean ± SEM.

